# Why the architecture of environmental fluctuation matters for fitness

**DOI:** 10.1101/2022.04.21.489085

**Authors:** John S. Park, Anja Felmy

## Abstract

The physical environment provides the very stage upon which the eco-evolutionary play unfolds. How fluctuations in the environment affect demographic fitness is thus central to selection predictions, life history analyses, and viability of populations. Treatment of fluctuating environments typically leverages the mathematics of random variability. However, environmental fluctuations in nature are almost always combinations of random and non-random components. For example, some fluctuations contain feedbacks which generate autocorrelation (*e.g.* disturbances such as floods, fires, and hurricanes), while others are driven by geophysical forces that create fixed cyclicality (*e.g.* seasonal, tidal, and diel). Despite theoretical developments, the consideration of non-random characteristics of fluctuations is still rare in empirical work on natural populations, mostly due to convention and partially due to difficulties in measuring and analyzing timeseries of environmental fluctuations. We show why non-randomness matters for fitness. Using a simple demographic model, we systematically compare four major categories of fluctuating environments: stochastic, positively autoregressive, negatively autoregressive, and periodic with error (“Noisy Clock”). The architectures of fluctuations influence the fitness of structured populations even when the modelled environments only differ in the timing of fluctuations, and not in their overall frequency. Importantly, we highlight two quantitative mechanisms through which fitness depends on fluctuation architecture—the consecutiveness of deviations from the environmental mean, and Jensen’s Inequality acting on nonlinear biological parameters—both relevant features in virtually all populations inhabiting variable environments. Our goal is to argue that non-random structures of environmental variability should be more seriously considered in empirical work. Such an endeavor would tap into the rich diversity of variable environments in nature to expand our understanding of the commensurate diversity of population dynamics.

## Introduction

Estimating population growth—or fitness—is a centerpiece of life history theory, evolutionary demography, and conservation biology. Reliable estimates of fitness have only become more important as climate change appears to increase variability in many systems, rendering the demographic consequences more varied and complex (Pike et al. 2004, Boyce et al. 2006, Davison et al. 2019). The potential for environmental variability to profoundly affect fitness outcomes is well-known. Yet, why and how types of environmental variability differ in their impact on fitness is less clear, especially in empirical work. Here we provide a synthesis of why and how the structure of environmental variability influences fitness. We first outline the history of conceptual developments, and then systematically compare the fitness consequences in four major categories, or ‘architectures’, of environmental variability. Lastly, we highlight two specific mechanisms that underlie fluctuations’ influence on fitness. We aim to spur a more careful consideration of the architecture of environmental variability in future empirical work.

There is a wealth of sophisticated theory on environmental impacts on demography and fitness. Cole (1954) first conceived of the fitness of life history strategies as directly linked to the strategies’ influence on the population growth rate of asexually reproducing organisms. Variations among genotypes in the age distribution of survival and reproduction shape the fitness landscapes of life histories (Gadgil and Bossert 1970, Stearns 1976), upon which natural selection can act. Evolutionary demography (Keyfitz 1968, Caswell 2001) leverages these concepts to analyze how the trajectories and sensitivities of population growth are influenced by population structures (*e.g.* compositions of ages, stages, or sizes) consisting of individuals on variable courses of life histories.

Early theoretical work such as above began with assumptions of constant environments for simplicity, and because it was necessary to lay the groundwork for comparing species in a systematic manner (Fisher 1958, MacArthur 1962). Developments since (*e.g.* Levins 1968, Schaffer 1974) have largely focused on extending analyses to include stochastic environment assumptions to reflect the complexities of nature (Lande et al. 2003, Tuljapurkar et al. 2003), and to spotlight the increasing environmental variability under climate change (Easterling et al. 2000, Drake 2005, Boyce et al. 2006, Morris et al. 2008, Lawson et al. 2015). Tuljapurkar *et al*. (2009) demonstrated why the expected optimal life histories differ between constant and stochastic environments, by decomposing the effects of life history speed, dispersion, and temporal correlation on fitness. Over time, ‘constant’ and ‘stochastic’ settled as the two dominant archetypical descriptions of environments, with the latter becoming the modern standard.

Stochasticity indeed offers a powerful suite of tools to deal with many kinds of variable environments, afforded by mature statistical theories of probability distributions. In short, a variable environment can typically be summarized by the first two moments of a distribution, *i.e.* the mean and variance, whose derivations lead to tractable analytical solutions for biological predictions in that environment (Lawson et al. 2015), *e.g.* bet-hedging strategies (Cohen 1966, Childs et al. 2010, Gremer and Venable 2014). Higher-order moments of stochastic environments can be used to analyze the effects of, *e.g.*, intra- and inter-annual variability and temporal autocorrelation on fitness (Tuljapurkar et al. 2009). There is substantial theoretical evidence that the ‘color’ of noise (negative to zero to positive autocorrelation) should influence fitness and extinction risk (Cohen 1966, Lande and Orzack 1988, Vasseur and Yodzis 2004, Halley 2005, Tuljapurkar and Haridas 2006, Reuman et al. 2008, Engen et al. 2013, Chevin et al. 2017). Overall, the subfield of ‘stochastic demography’ uses stochastic environmental formulation at its core to ask ecological and evolutionary questions (Tuljapurkar 1990, Fieberg and Ellner 2001, Engen et al. 2009, Lande et al. 2017, Chevin et al. 2017, Schreiber and Moore 2018, Crewe et al. 2018, Davison et al. 2019).

Mathematically, stochasticity is specific: it describes a process that can be captured with a random probability distribution. However, many real categories of environmental fluctuations have non-random components. For example, fundamental geophysical fixtures such as the rotation of Earth (governing diel cycles), its axial tilt and revolution around the Sun (seasonal cycles), and the revolution of the Moon around Earth (tidal cycles) set periodic oscillations in most terrestrial and aquatic ecosystems. Analogous to driven harmonic oscillators in physics, such environments are governed by fixed (at least on ecological and evolutionary time scales) external drivers of fluctuations. Distinct from stochastic variability, geophysically-driven fluctuations are thus not emergent products of the internal ecological and evolutionary dynamics of the system such as in population cycles or ecological succession.

The particular sequence of environmental fluctuations is biologically important, over and above the overall amount of environmental variability. Stearns (1976, Fig. 7) outlined the possible outcomes of environmental fluctuation frequency and predictability for the age distribution of reproduction in a population. Later, Tuljapurkar (1985) analyzed how the relative scaling between the period of environmental cycles and generation time influences population growth rate. Recent work demonstrated that environmental periodicity may itself be a selective force that determines both optimal (Park 2019) and individual variation in life histories (Park and Wootton 2021).

Empirically speaking, climate change is rapidly perturbing seasonality and fluctuation regimes, with far-reaching consequences such as phenological shifts (Walther et al. 2002, Westerling et al. 2006, Marlon et al. 2009). Importantly, the mean and variance of environmental fluctuations can change independently in disturbance regimes (Lytle 2001, Lytle et al. 2008, Denny et al. 2009), which is a growing focus in climate change biology (Jentsch et al. 2007, IPCC 2012, Thornton et al. 2014). Thus, teasing apart the non-random and random components that constitute environmental variability—instead of collapsing all environmental variability to stochasticity—is crucial not only for more accurate predictions of population growth (Tuljapurkar and Haridas 2006), but for systematizing the variety of causal drivers of population dynamics and life history selection.

We focus on four broadly representative architectures of fluctuating environments and use a simple demographic model to show the architectures’ contrasting effects on demographic fitness. The four environments solely differ in the exact timing sequence of events. Events here represent ecological disturbances (Lytle 2001) such as fires, floods, hurricanes, avalanches, droughts, and heatwaves, or, in seasonal systems, onsets of winter that can largely impede biological activity (Inouye 2008). A particularly novel consideration is the fourth category of fluctuations: inherently periodic environments such as seasonal or tidal ones. These externally driven cycles with fixed periodicity are fundamentally different in their memory process compared to internally produced cycles (*e.g.* autocorrelated): in the former, environmental states gravitate to an external schedule, whereas in the latter, environments depend on former states of themselves. We then proceed to identify two general quantitative mechanisms that cause divergences in fitness estimates among all four categories. The first mechanism involves the pattern of consecutive deviations from the environmental mean. The second involves Jensen’s Inequality acting on nonlinear life history traits, or vital rates; the prevalence of such nonlinearity in nature is likely universal. Both are very general properties of populations in variable environments. We do not claim that these are the only mechanisms at play, nor that our list of fluctuation categories is exhaustive. We also do not generate new theory. Instead, our goal is to show how the two mechanisms influence fitness even in simple situations, and to systematically highlight the role of non-random variability beyond the conventional archetype of random variability. Finally, we discuss the general relevance of our findings and identify key extensions and empirical tests that will be important moving forward.

### Four categories of fluctuating environments

We start by setting up illustrative examples of four major categories of fluctuating environments (Fig. 1): stochastic (white noise), positively autoregressive (red noise), negatively autoregressive (blue noise), and “Noisy Clock” (fundamentally periodic, but with error). For all four categories, we focus on the parameter *τ*_*i*_which is the *interval* between events. Again, these events represent ecological disturbances such as fires, or the onset of winter in seasonal systems with the assumption that winters are harsh enough to perturb or halt biological activity. In seasonal systems, note that celestial seasons are perfectly periodic, but the meteorological onset of winter (*e.g.* temperature, frost) and arrival of spring are variable, making the intervening warm season (the interval) also variable in length (‘climatological growing seasons’, *sensu* Linderholm 2006). We derive the mean and variance of the distribution of intervals between events from each category to make them exactly analytically equivalent, before comparing the fitness consequences in environments characterized by the different fluctuations.

**Figure 1.**
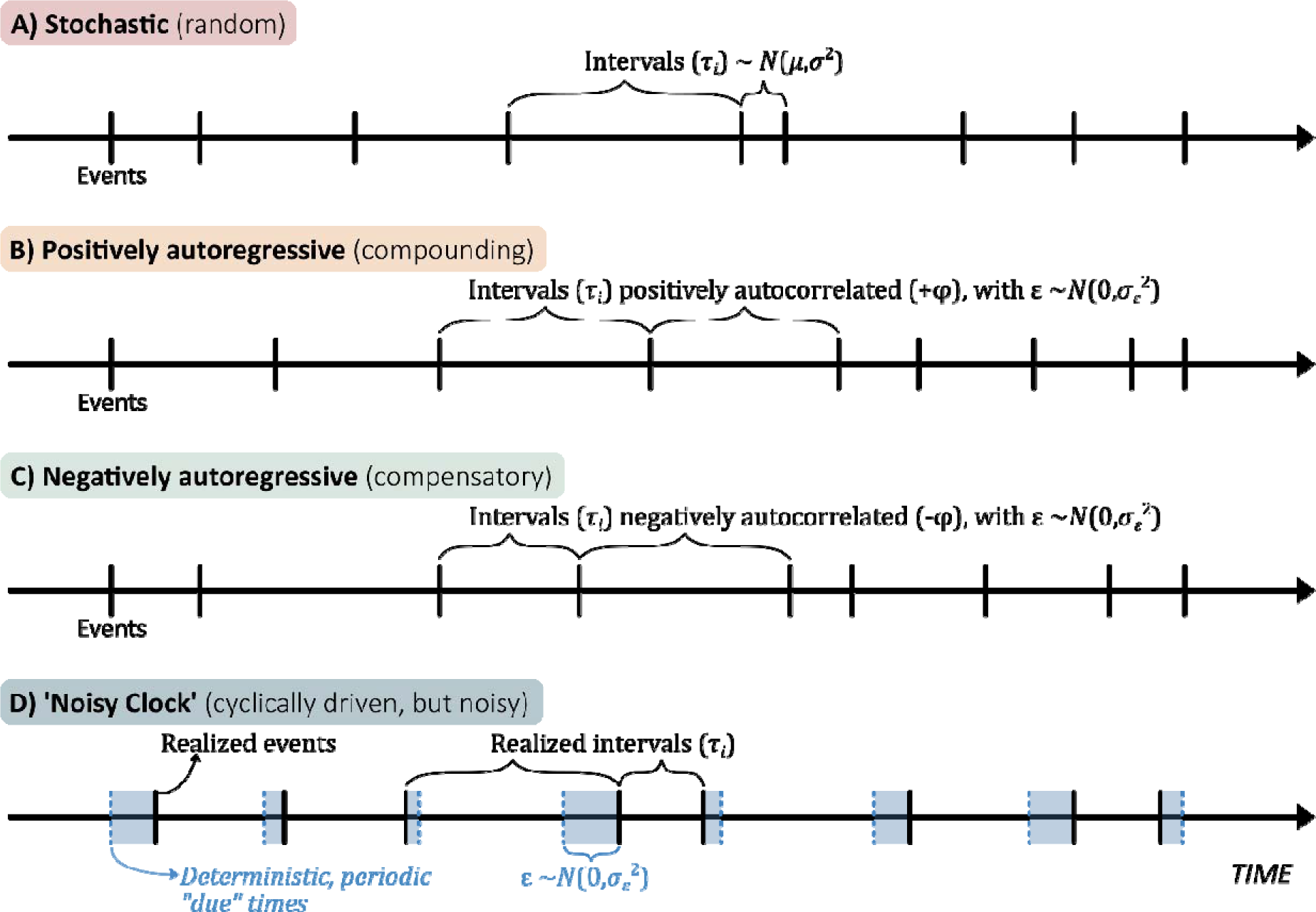
Four categories of fluctuating environments. Each fluctuation architecture is simulated by generating a sequence of event intervals on a timeline. Events, denoted by the tick marks, may represent ecological disturbances or winters that bring harsh conditions for biological activity. A) A stochastic environment is generated by drawing a sequence of intervals from a random (Normal) probability distribution. B) A positively autoregressive environment is generated with positive linear dependency of subsequent intervals on immediately preceding ones (an AR(1) process), with random error; C) the negatively autoregressive environment is only different from B in that the linear dependency on the past interval is negative. D) A Noisy Clock environment assumes a series of unseen “due times” that are perfectly periodic (dashed blue tick marks), but the actual events occur as a result of error around those due times (blue shaded regions); the gaps between realized events form the sequence of realized intervals.

#### Category I: Stochastic (random), without feedback

We generate a fluctuating environment with a random sequence of intervals *τ*_*i*_ between environmental disturbance events:

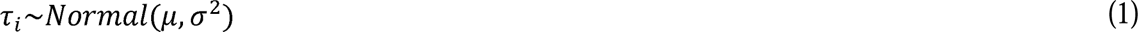

where *μ* is the desired interval mean and *σ*^2^ the variance. We do this instead of generating a series of events via a Poisson process, which is a more common way of generating random intervals on a timeline, to make this case comparable with the other three cases with respect to their error distributions and because below we will manipulate the amount of noise (*σ*^2^) while holding the mean interval constant (which is not possible with Poisson distributions because their mean and variance are equal). While using a normal distribution does not theoretically ensure negative values cannot be drawn—which would be nonsensical for interval length—for simplicity we have set the mean (=100) and variances (ranging from 0 to 300) of our simulations such that the probability of a negative value is exceedingly low (< 3.9 × 10^−9^); we then confirmed that negative values were in fact not drawn in our simulations.

Importantly, note that this manner of generating a fluctuating environment does not consider the source or generative nature of the fluctuations, nor the length of preceding intervals (*i.e.* there is no memory of past fluctuations), but only assumes the long-run statistical distribution of variability (Fig. 1A). While measured environmental variability may in reality always have a physical source, and all systems may have some memory process to varying degrees, some systems are so complex and noisy that they are well approximated by pure randomness. Random environments may be experienced, for example, by mice living in dryland ecosystems with variable resource availability (Noble et al. 2019), by weevil species using strongly fluctuating oak acorns as egg-laying sites (Pélisson et al. 2012), and by annual herbs growing in or near vernal pools created by stochastic winter precipitation events (Emery and La Rosa 2019).

#### Category II: Positively autoregressive

We generate a positively autocorrelated environment with a simple AR(1) process:

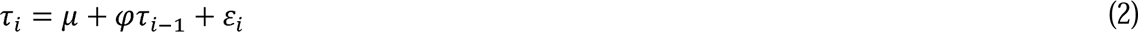

where *μ* is the imposed mean interval between environmental disturbances, and the interval *τ*_*i*_is positively correlated by strength of autocorrelation *φ* with the interval immediately preceding it (*τ*_*i*−1_). The i.i.d. (independent identically distributed) noise *ε*_*i*_ is drawn from 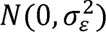. Positively autocorrelated environments are characterized by a ‘compounding’ behavior where a longer or shorter interval than the mean is likely to beget another longer or shorter interval in the immediate future, respectively (Fig. 1B). Examples of positively autocorrelated fluctuations in nature include some fire regimes (Caswell and Kaye 2001), oceanic temperatures (Steele 1985, Halley 2005), and flood regimes (Sabo and Post 2008).

#### Category III: Negatively autoregressive

The negative counterpart to the positively autoregressive environment is one with a negative autoregressive term (−*φ*) and all other terms identical to Eq. 2:

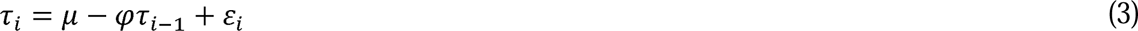

Negatively autocorrelated environments are characterized by a ‘compensating’ or self-correcting behavior where an interval between disturbances that is longer or shorter than the mean is likely to be counterbalanced with a shorter or longer interval in the immediate future, respectively (Fig. 1C). Real examples of negatively autocorrelated environments in nature, though much more rarely considered than positively autocorrelated ones (Metcalf and Koons 2007), include terrestrial plant systems with delayed density-dependence driven by nutrient cycling (Gonzalez-Andujar et al. 2006), and tree masting (Hacket-Pain and Bogdziewicz 2021).

#### Category IV: ‘Noisy Clock’

The last and least explicitly considered category of fluctuations, but one that is ubiquitous in nature, includes environments with a fundamental external driver of cyclicality, whose deterministic periodicity is not affected by the realized noise. We henceforth refer to this case as the ‘Noisy Clock’ to describe the clockwork (deterministic) nature of fluctuations that underlies the noise. Here, we first punctuate the timeline with a series of deterministic ‘due’ times of events much like the celestial periodicity of Earth’s revolution that determines the boundaries of the annual cycle. Intervals between disturbances are not generated by counting time from the preceding disturbance events, but instead, imperfectly timed disturbances events occur around each deterministic due time with i.i.d. noise (Fig. 1D). Then the intervals are the gaps between subsequent realized events:

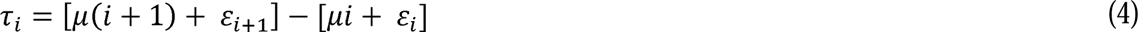

where 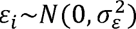 and 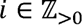, meaning clocked due times are positive integer multiples of *μ*, the same mean interval as in categories I, II, and III. Counting time from preceding events (which is the case for categories I, II, and III) as opposed to in reference to an external clock would be an unrealistic way to model fundamentally cyclical systems such as seasonal ones. For instance, if one were to model a winter event (*e.g.* first snowfall), but stochastically modelled the length of intervening summer seasons with no regard to due times, then after some additive sequence of stochastic lengths it would be possible to have winters starting in July when lined up against a deterministic calendar. This would be unrealistic under any reasonable geophysical stationarity assumptions. Adding error to due times has the effect of gravitating noisy events back to realistic due times set by the underlying forces of harmonic oscillations in the environment. Real fluctuations with clocked due times driven by geophysical forces—but with error in the measured proxy variables—are exemplified by a myriad of diel (*e.g.* hunting behavior of African wild dogs; Nouvellet *et al*. 2012), tidal (*e.g.* foraging of intertidal snails; Hayford *et al*. 2018), and seasonal environments (*e.g.* advancement of egg-laying with warming springs in numerous bird species; Dunn & Møller 2014). Despite widespread attention to seasonal environments, particularly with respect to global phenological shifts (Walther et al. 2002), the theoretical foundation of demographic dynamics and life history evolution in cyclical environments is not yet well understood (Lande et al. 2017).

### Comparing apples to apples: the same normal distribution

Probability distributions that generate timeseries can be described by their moments, the first of which are the man, variance, skewness, and kurtosis. Conventionally, a fluctuating environment under study is analyzed or modelled using the first two moments – the mean and variance of event timing or of some continuous parameter (*e.g.* temperature). However, fluctuating environments often also differ in their higher-order structures such as autocorrelation in the timing of environmental disturbances and intervals between them. Here, we additionally consider a structure that is neither purely stochastic nor simply autocorrelated, but one that has a fundamental deterministic periodicity (Noisy Clock). To compare them systematically, we first derive input parameters that would make the mean and variance of the four fluctuating environments (Eqs. 1-4) analytically equivalent (Table 1; Fig. S1). We set mean interval (*μ*) to equal 100 time steps, and equivalently vary the variance across the four environments to see its effect on fitness. Then we explore why even given this equivalency, fitness estimates diverge among the environments. Our goal is to demonstrate that the conventional assumption that any fluctuating environment is sufficiently well-characterized as a random environment with mean and variance of the fluctuation—while justified in some cases—can subtly, but consistently, lead to inaccurate estimates of fitness.

**Table 1.** Expressions of the expected value and variance of the four fluctuations, and the input parameters in terms of the desired sample moments (terms with subscript *s*) that will force *τ*_*i*_’s of all four fluctuations to be equivalently distributed as the normal distribution of category I.

| | Fluctuation | Expected Value ($E[\tau]$) | Variance ($Var[\tau]$) | Input parameters in terms of $\mu_s$ and $\sigma_s^2$ |
| --- | --- | --- | --- | --- |
| I | $\tau_i \sim Normal(\mu, \sigma^2)$ | $\mu$ | $\sigma^2$ | $\mu = \mu_s;$<br>$\sigma^2 = \sigma_s^2$ |
| II | $\tau_i = \mu + \varphi\tau_{i-1} + \varepsilon_i,$<br>where $\varepsilon_i \sim N(0, \sigma_\varepsilon^2)$ | $\mu$ | $\frac{\sigma_\varepsilon^2}{1 - \varphi^2}$ | $\mu = \mu_s;$<br>$\sigma_\varepsilon^2 = \sigma_s^2(1 - \varphi^2)$ |
| III | $\tau_i = \mu - \varphi\tau_{i-1} + \varepsilon_i,$<br>where $\varepsilon_i \sim N(0, \sigma_\varepsilon^2)$ | $\mu$ | $\frac{\sigma_\varepsilon^2}{1 - \varphi^2}$ | $\mu = \mu_s;$<br>$\sigma_\varepsilon^2 = \sigma_s^2(1 - \varphi^2)$ |
| IV | $\tau_i = [\mu(i + 1) + \varepsilon_{i+1}] - [\mu i + \varepsilon_i],$<br>where $\varepsilon_i \sim N(0, \sigma_\varepsilon^2)$ and $i \in \mathbb{Z}_{>0}$ | $\mu$ | $2 \times \sigma_\varepsilon^2$ | $\mu = \mu_s;$<br>$\sigma_\varepsilon^2 = \frac{1}{2}\sigma_s^2$ |

### A simple demographic model in fluctuating environments

Upon the environmental fluctuation backgrounds (Fig. 2A), we model a simple population that proceeds through time. There are 2 time indices: *t* denotes the discrete time steps, here imagined as days (but scalable depending on system), and *τ*_*i*_ indexes the interval lengths (measured in time steps) between subsequent disturbance events, the sequence of which is determined by the categories of fluctuation (Eqs. 1-4). *J*_*t*_ and *A*_*t*_ are numbers of juveniles and adults at time *t* respectively, *f* is the per capita rate of reproduction, ℓ is the rate of juveniles maturing into adults, and *S*_*J*_ and *S*_*A*_ are rates of juvenile and adult survival. The population dynamic is then given by the system of equations:

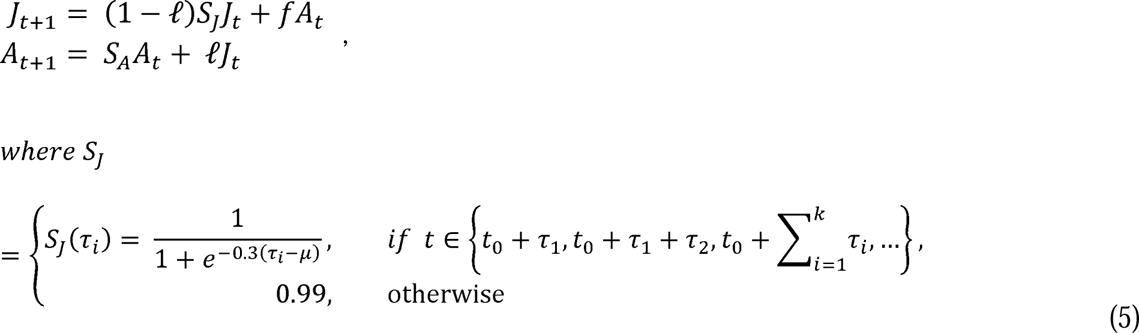

**Figure 2.**
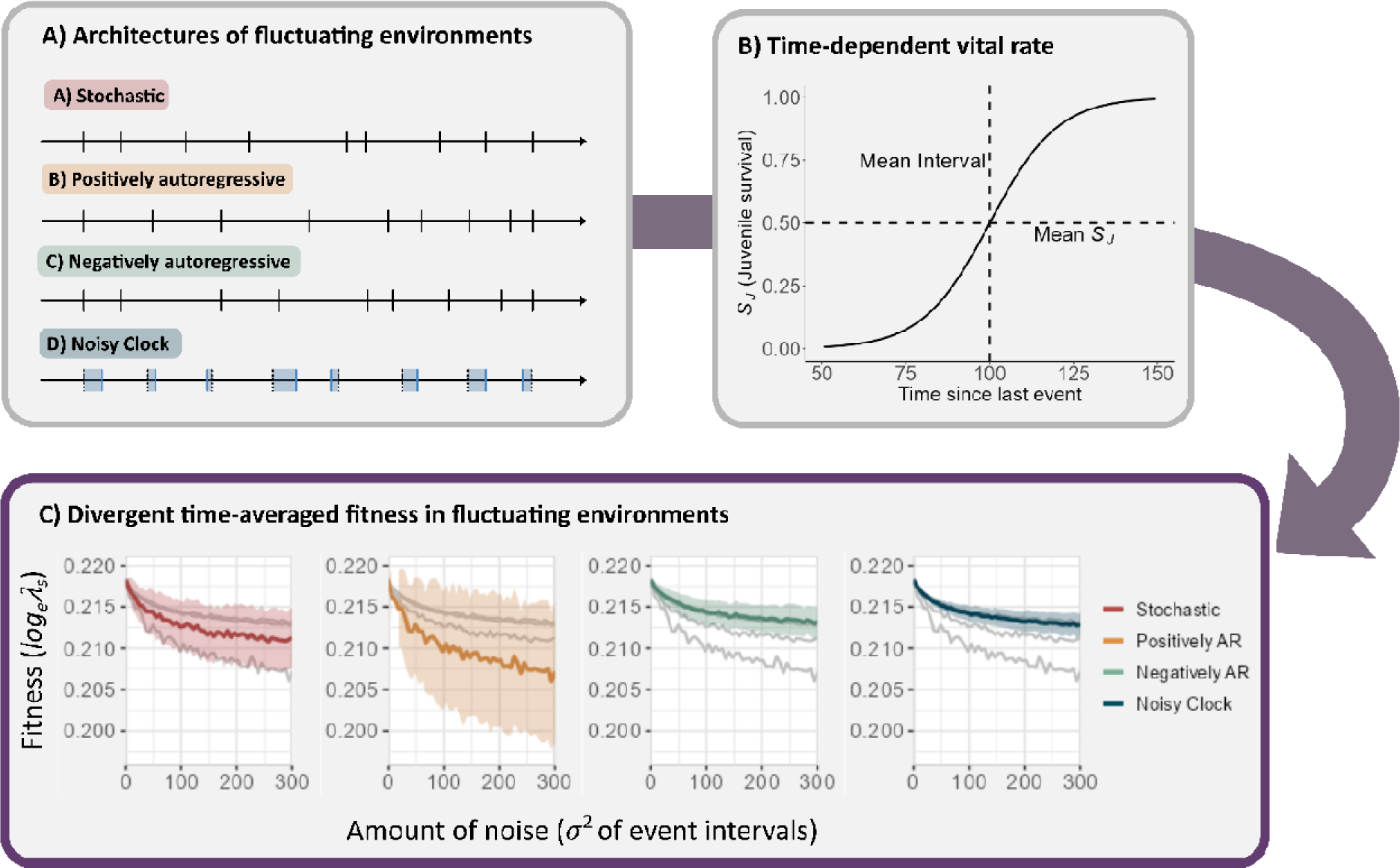
The interaction of environmental fluctuation architecture and nonlinearity in a vital rate creates divergence in fitness estimates. A) Diagrammatic representation of the four architectures of fluctuation, through which the demographic dynamics proceed. B) Juvenile survival rate (*S*_*J*_) increases nonlinearly as a function of the length of the time passed since the previous disturbance events. Dashed lines indicate the means of each axis. C) When passed through B, the different architectures of A result in divergent fitness outcomes. Panels are split for better visualization. Each colored line shows the mean time-averaged fitness (*log*_*e*_*λ*_*s*_) over 500 iterations with increasing *σ*^2^, after 500 event intervals (500 × *μ* = 50,000 *time steps*). Bands show ±1 *sd* over the iterations. Grey lines in each panel show the mean lines of the other 3 environments. Fitness is driven apart further as *σ*^2^ (amount of noise, or unpredictability, in the environment) increases.

We focus on juvenile survival *S*_*J*_ as the vital rate that actively interacts with the disturbances that arrive intermittently into the system. This is because in many natural populations of plants or animals juvenile mortality is known to be especially sensitive to harsh environmental conditions such as disturbances, and particularly to how frequently they arrive (Gaillard et al. 1998, 2000, Higgins et al. 2000, Lytle 2002, McAdam and Boutin 2003, Staver et al. 2009, Park 2019). To reflect this in Eq. 5, juvenile survival *S*_*J*_ is determined by a logistic response function *S*_*J*_(*τ*_*i*_) (Fig. 2B) when disturbance occurs (*i.e.* when time step *t* is an element of the predetermined sequence of disturbance timings, which is the cumulative sum of successive interval lengths 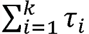, where *k* is the total number of disturbance intervals simulated), and is 0.99 otherwise (in between disturbances). The shape parameter 0.3 in the denominator is the steepness of the logistic curve, the inflection point is at *S*_*J*_ = 0.5, and the difference *τ*_*i*_ − *μ* determines how far above or below *S*_*J*_(*τ*_*i*_) is from mean *S*_*J*_. Verbally, this means that if two disturbance events happen in quick succession, juvenile survival will be low, and if a long time passes between events, it will be high, in a nonlinear fashion. Biologically, this function signifies the time needed for juveniles to sufficiently develop, grow, or learn to endure the next impending perturbation in the environment. Though not always logistic, nonlinearity in vital rate response functions in between perturbations in the environment is likely common (Caswell 2008, Childs et al. 2010, Sletvold et al. 2013, Taylor et al. 2014, Ehrlén et al. 2016, Shriver 2016). For example, Morris et al. (2006) showed that, across five species of perennial plants, vital rates including seedling survival (equivalent to *S*_*J*_here), survival at other stages, growth, germination, and seed production (equivalent to *f* here) changed nonlinearly as a function of time since last hurricane or fire disturbance. Similarly, Bürkli & Jokela (2017) showed that embryonic development success (early-stage juvenile survival) of a freshwater snail changed nonlinearly across the species’ reproductive season. The exact shape of vital rate response functions is another key point to be tailored for system-specific inquiry, whether with empirical measurements or as realistic assumptions built into models.

The model is otherwise kept simple to focus on how the different categories of environmental fluctuation affect fitness through just one vital rate. Time-step values for all parameters are biologically realistic for daily rates (*e.g.* Least Killifish, a small live-bearing freshwater fish (Felmy et al. 2021)): we hold adult survival *S*_*A*_ constant at 0.99 regardless of disturbance, and maturation rate ℓ at 0.01 (thus birth-to-birth generation time ≈ mean disturbance cycle length,

100). Rate of reproduction *f* is logistically density-dependent, where 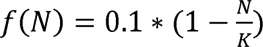 where N is total population size at *t* and carrying capacity K=1000. Structural complexity such as the number of developmental stages (*i.e.* the number of equations in the system, or the number of vital rate terms), or whether multiple vital rates interact with disturbance frequency, would be an obvious area of customization for system-specific inquiry. The downstream results remain qualitatively unchanged when *f*, ℓ, and *S*_*A*_ are allowed to vary within a reasonable range (Fig. S2). As we will argue, the key mechanisms should influence fitness regardless of demographic nuance.

### Time-averaged fitness

In each of the four architectures of environmental fluctuations, we ask what the long-run fitness of a life history (the set of vital rate parameters defined above) would be. For fitness, we use the standard

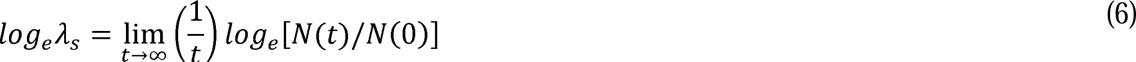

which gives the long-run growth rate in fluctuating environments (Tuljapurkar 1990) where *N*(*t*) is the total population size at time *t*, including both juvenile and adult stages. This metric can be computed from numerical simulations over a long time course of the environment (Caswell 2001), and robustly approximates fitness even under considerably large fluctuations (Lande et al. 2003, Tuljapurkar et al. 2003, 2009). We simulate demographic dynamics (Eq. 5) in each fluctuating environment (Fig. 1) for 500 event intervals with mean interval (*μ*) = 100 time steps (∼50,000 time steps), calculate *log*_*e*_*λ*_*s*_, and repeat these calculations over 500 iterations. Lastly, we compute all of the above across a spectrum of variance in event intervals (0 to 300), to assess how the amount of fluctuation noise affects the divergence of fitness estimates.

### Fitness divergence across categories of fluctuating environments

Our results show that fitness estimates of the same population model differ substantially among the four architectures of fluctuating environments (Fig. 2C), despite the analytical equivalency of the environments’ distributions of intervals between disturbance events (Table 1; Fig. S1). Across the spectrum of environmental variance, *log*_*e*_*λ*_*s*_ was overall highest for the negatively autoregressive environment, slightly lower for the cyclically driven but noisy environment (Noisy Clock), still lower for the stochastic environment without autocorrelation, and by far the lowest for the positively autoregressive environment. Importantly, all above differences in fitness among the four environments are amplified as *σ*^2^ is increased.

### Mechanisms of fitness divergence: Consecutiveness of deviations and Jensen’s Inequality

The central question that arises from Fig. 2C is why fitness estimates diverge between the four architectures of fluctuating environments, despite the environments’ distributions of event intervals being forced to be analytically equivalent (Table 1). There are two mechanisms that work in concert to drive the divergence: (1) *Consecutiveness of deviations from the environmental mean,* and (2) *Jensen’s Inequality in vital rate fluctuations*.

#### (1) Consecutiveness of deviations from the environmental mean

In fluctuating environments with a mean interval between environmental disturbances, deviations to one side of the mean interval (*i.e.* below or above) can have short consecutive runs (*e.g.* flipping equally and quickly between below and above the mean) or long consecutive runs (*e.g.* many deviations below or above the mean in a row) even if the long-term overall distribution of deviations is normal. The four categories of environmental fluctuations have different levels of compounding or compensatory behavior of fluctuations, creating different distributions of consecutiveness (Fig. 3A). Negatively autoregressive and Noisy Clock environments both tend to self-correct deviations back to the mean, driven by the −*φ* in the former, and by the deterministic ‘due times’ in the latter. Thus, these environments have short tails in their distributions of consecutive deviation length. In contrast, the positively autoregressive environment by definition repeats deviations to one side of the mean, resulting in a long tail. The stochastic environment exhibits an intermediate exponential distribution of consecutive deviations.

**Figure 3.**
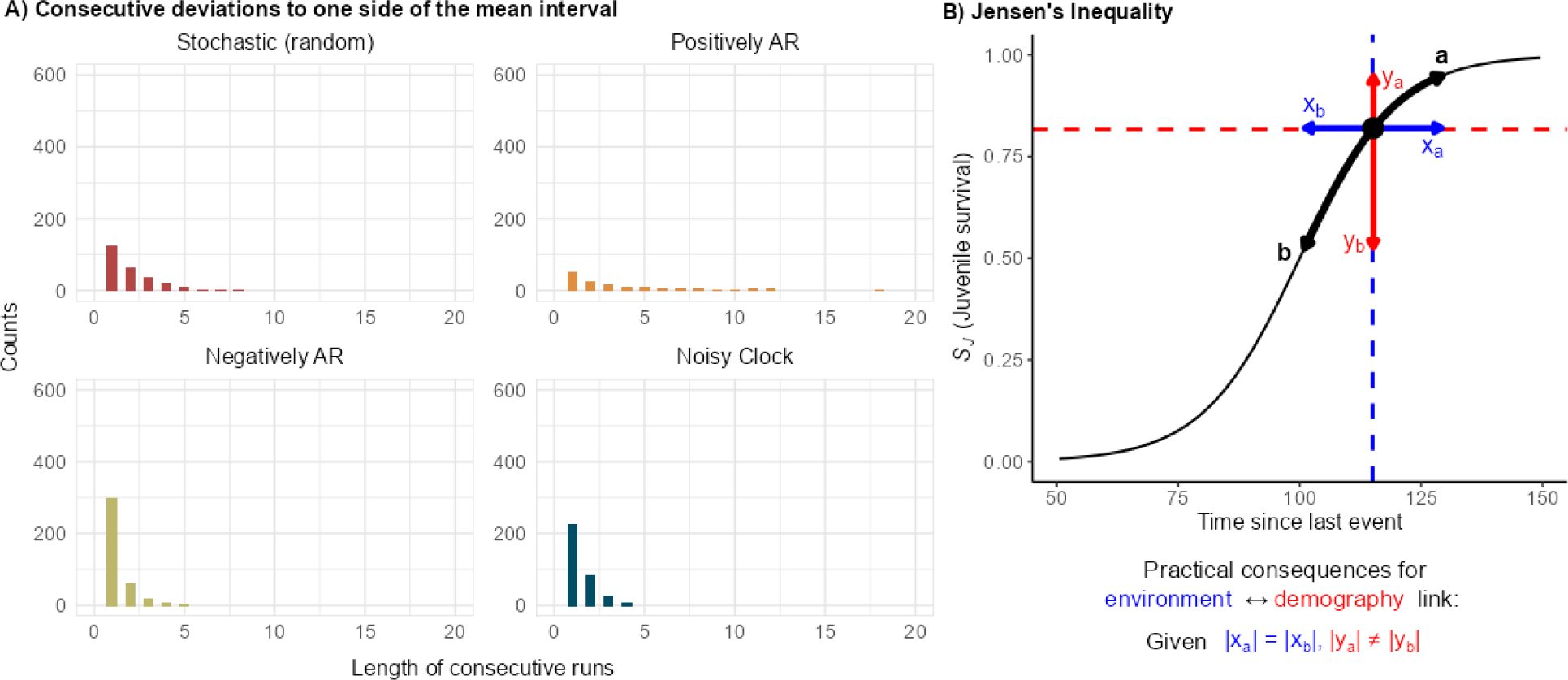
Mechanisms of fitness divergence between the architectures of fluctuating environments. A) Despite analytically equivalent distributions of event intervals (Table 1), environments have very different distributions of consecutive runs to one side of (below or above) the mean interval depending on the nature of the generating function. Negatively autoregressive and Noisy Clock environments inherently exhibit compensatory or self-correcting behavior, resulting in shorter tails in the consecutive run distribution. The opposite is true of the positively autoregressive case. The stochastic environment exhibits an intermediate exponential distribution. B) Jensen’s Inequality exists in any convex/cave fluctuating variable, and introduces an asymmetry between environmental and demographic fluctuations. Nonlinear responses in vital rates as a function of interval length between ecological events such as disturbances are likely common.

Why does the distribution shape of consecutive deviations matter for fitness, if over time, there is an identical symmetrical accumulation of negative and positive deviations (*i.e.* normal distribution of intervals between disturbances; Table 1 and Fig. S1)? Long-run fitness *log*_*e*_*λ*_*s*_is a multiplicative, or geometric, fitness measure. Consecutive runs of positive deviations of a vital rate (*e.g.* ‘good’ years) will never evenly compensate for equally long consecutive runs of negative deviations (*e.g.* ‘bad’ years) with respect to population growth (Haccou and Vatutin 2003, Cuddington and Hastings 2016). To illustrate this point, imagine an extreme scenario in which a population of organisms relies on fog for moisture and can only survive for one day without it. In environment A, a sunny day occurs on average every 100 days and is always followed by a foggy one. Hence, the population persists. But in environment B, sunny days always come in consecutive pairs, occurring every 200 days on average. Although sunny days are equally common in both environments over the long run, a population inhabiting environment B will soon be extinct. Therefore, in structured populations, the distribution of the lengths of consecutive deviations should influence long-run fitness (longer tail = lower fitness).

#### (2) Jensen’s Inequality in vital rate fluctuations

Fluctuations below and above the mean disturbance interval also have asymmetrical consequences for the corresponding nonlinear demographic parameter; here that parameter is juvenile survival *S*_*J*_, which has a direct effect on fitness *log*_*e*_*λ*_*s*_. This asymmetry is referred to as Jensen’s Inequality (Jensen 1906, Ruel and Ayres 1999), which states that for a convex or concave function *f*(*x*), *E*[*f*(*x*)] ≤ *or* ≥ *f*(*E*[*x*]). Verbally, this means that symmetrical fluctuation of a variable *x* about its mean along the x-axis does not translate equivalently to symmetrical fluctuation about its mean along the y-axis. If we take fluctuation along the x-axis as that of the environment, and fluctuation along the y-axis as that of demography, Jensen’s Inequality introduces a break in the environment-demography link in fluctuating environments (Fig. 3B).

The direction of that asymmetry (*i.e.* sign of the inequality) and its magnitude depend on the convex/cavity of the region of the nonlinear function where the mean lies. We demonstrate the consequence of this dependency by manipulating the convex/cavity of the nonlinear demographic parameter *S*_*J*_around the mean. Consequently, the mean *S*_*J*_, which was 0.5 in the prior demonstration (Fig. 2C), will now equal 0.1, 0.3, 0.5, 0.7, or 0.9 (Fig. 4A). When mean juvenile survival is low and its response function convex around the mean (Fig. 4A, panels towards the left), fitness increases with increasing environmental variability *σ*^2^ (corresponding panels in Fig. 4B). This is because deviations to the right of the mean interval over-compensate for deviations to the left in terms of survival *S*_*J*_and thus boosts long-term growth of the population. Conversely, when mean juvenile survival is high and its response function concave around the mean (Fig. 4A, panels towards the right), fitness curves are pulled down with increasing *σ*^2^. The reverse explanation of the former explains this behavior. Note that, when mean interval is exactly at the inflection point of the response function (*S*_*J*_ = 0.5; Fig. 4A center panel), Jensen’s Inequality does not come into play, as deviations to either side of the mean interval equally compensate one another in terms of their effects on survival. Regardless of the convex/cavity of the response function of juvenile survival, as *σ*^2^increases fitness curves in all four environments begin to stabilize; this occurs because with higher environmental variance, fluctuations begin to sample from wider breadths of the logistic response function which has a combination of convex and concave regions, weakening the role of Jensen’s Inequality. Nevertheless, the relative ordering of fitness across fluctuation architectures remains stable (Fig. 4B, all panels), demonstrating the role that the consecutiveness of deviations from the environmental mean plays in setting and maintaining the divergence of fitness between fluctuation architectures.

**Figure 4.**
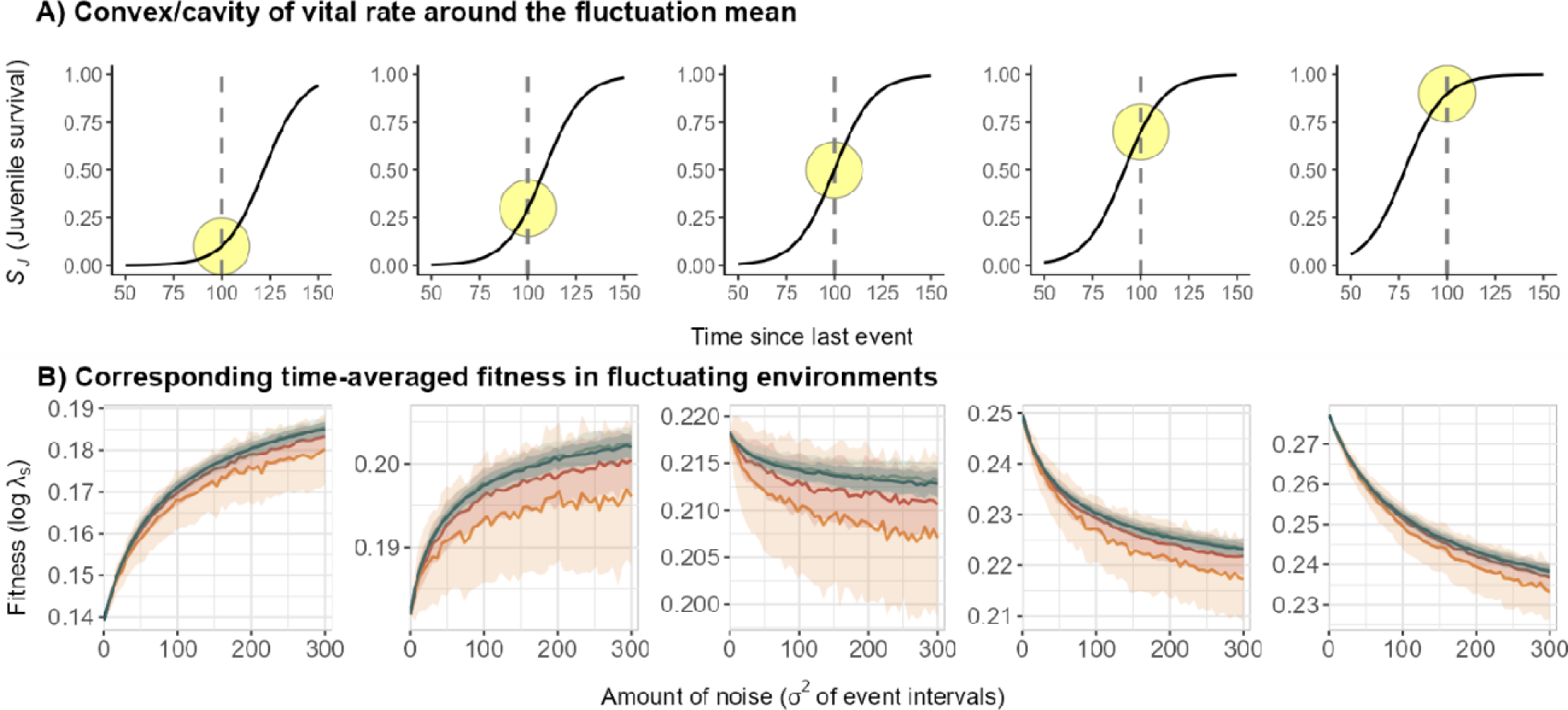
Demonstration of the effect of Jensen’s Inequality on fitness divergence across architectures of environmental fluctuation. A) We shift the response function of juvenile survival *S*_*J*_, and thus mean *S*_*J*_ (*S*_*J*_ at mean event interval), to manipulate the nonlinear shape around the mean event interval (dashed line), which is held constant. Going from left to right, the response function is more convex to more concave around the mean interval, highlighted in the yellow circles. This flips the sign of Jensen’s Inequality, so that the overall effect of environmental fluctuations on fitness would generally become more positive to negative. B) The corresponding fitness plots demonstrate the sign flip in the effect of Jensen’s Inequality as a result of the convex/cavity around the mean. Colored lines are means over all iterations, and bands are ± 1 sd. Colors correspond to the four architectures in previous figures. The middle panel is equivalent to a composite of Fig. 2C.

### Summary and Future Outlook

Non-random temporal environmental variability abounds in nature, from autocorrelated fluctuations to externally driven cycles (Sabo and Post 2008, Dunn and Møller 2014, Hacket-Pain and Bogdziewicz 2021). Yet the rich diversity of non-random fluctuations is often subsumed under the convention of stochasticity (described by random probability distributions) in ecological and evolutionary models, for its alluring mathematical properties and ‘good enough’ approximations of nature. We have shown that for fitness, the architecture of non-random components of variability matters, even when long-term environmental variabilities are distributed equivalently.

In itself, incorporating non-randomness in environmental models is not new to many researchers working on life history theory or demography. Developments in periodic matrix models (Caswell and Trevisan 1994), the seasonal forcing of disease dynamics (Altizer et al. 2006), bet-hedging strategies (Venable 2007), and the effect of temporal autocorrelation on life history evolution and population persistence (Pike et al. 2004, Metcalf and Koons 2007, Tuljapurkar et al. 2009) are some examples of considering non-random fluctuations. The novelty of our results is two-fold: we systematically show that random and non-random fluctuations lead to consistent differences in demographic fitness; then, we highlight two simple but general mechanisms that drive this divergence. These mechanisms are prevalent, and their treatment should open new doors for investigating demography and selection dynamics in the diverse—and changing—fluctuation architectures in nature.

Our results corroborate a message that has repeatedly emerged out of the theoretical literature of colored noise and extinction risk, namely that “red” environments (positively autocorrelated) increase extinction risk while “blue” environments (negatively autocorrelated) do the opposite (Lawton 1988, Halley and Kunin 1999, van de Pol et al. 2011, Halley et al. 2018). Although we focused on long-run fitness as opposed to extinction risk *per se,* the link to population persistence is obvious: in positively autocorrelated environments, the long-run growth rate was not only lowest by far, but also showed the highest amount of variance (Fig. 2C), increasing the likelihood of a population crash proving fatal. In contrast, negatively autocorrelated fluctuations led to the highest, most constant long-run fitness (Fig. 2C). Importantly, we further distinguish autocorrelated environments from Noisy Clock environments (*e.g.* seasonal, tidal, and diel). Autocorrelated environments rely on the memory of past events, whereas Noisy Clock environments, in their simplest form, are memoryless and instead anchored to an external clock such as Earth’s spin and revolution around the sun. This subtle difference—invisible through distributions of intervals between environmental disturbances—is visible in the distribution of consecutive deviations from the fluctuation mean (Fig. 3A), and thus long-run fitness. In our simulations, fitness in Noisy Clock environments consistently fell between that of red and blue environments (Figs. 2C & 4B), suggesting that modelling a seasonal environment simply as an autocorrelated one might lead to erroneous calculations of fitness. This result further suggests that Noisy Clock environments, due to their locking onto an external clock, stabilize population growth compared to other types of fluctuations; mathematical proof of this notion would be interesting and broadly relevant as Noisy Clock-like environments are ubiquitous.

In reality, environmental fluctuations will seldom neatly fall into only one of the architectural categories simulated here. For example, a coastal system that is strongly perturbed by periodic tidal disturbances will also be influenced by seasonal fluctuations (Denny et al. 2009). How resonances of non-random fluctuations constructively or destructively interfere to produce a variety of fluctuation superpositions in different ecosystems might explain a wider breadth of observed life history diversity and population trajectories. Even if an environment is predominantly driven by one type of fluctuation architecture, the equivalence of the interval distributions as we showed here can make it difficult to place a given environment into one fluctuation category. Empiricists could consider plotting the distribution of consecutive deviations from the measured mean fluctuation interval, as we did in Fig. 3A. The shape of this distribution, compared against the null of a distribution of consecutive deviations in a purely random environment (Fig. 3A, “stochastic”) might give an indication of whether an environment is fluctuating positively autoregressively (long tail in the distribution of consecutive deviations), or negatively autoregressively / in a Noisy Clock fashion (short tail). Climate scientists and population ecologists regularly use spectral analyses to detect dominant periodicities and lag effects in time series of disturbance events or continuous environmental variables (Priestley 1981, Chatfield 2003, Sabo and Post 2008), though often to remove temporal autocorrelation so that long-term trends can be detected (Pyper and Peterman 1998, Brown et al. 2011). We argue that consideration of such methods should become more common for life history and evolutionary demography analyses in nature. Our demonstrations suggest that interactions between the architecture of environmental fluctuation and nonlinear biological vital rates (*e.g.* survival, growth, reproduction) will alter fitness estimates in a broad range of situations.

Notably, these fitness differences arise from environmental sensitivity in just one vital rate— juvenile survival—while adult survival and fecundity were modelled as environment-independent. For generality, we strived to simplify our life history model, but its degree of complexity is likely another fruitful avenue of expansion and customization for specific empirical tests. For instance, many species have more than two distinct life stages, for example amphibians (*e.g.* egg, larva, adult) and insects (*e.g.* egg, nymph, pupa, adult). The number and interconnectedness of stages via growth, survival, and reproduction might have other complex and interesting influences on how fitness differs between fluctuation architectures. The nonlinear response function of a vital rate can take many different shapes depending on the proximate developmental, physiological, or behavioral mechanisms, from quadratic to self-excitatory threshold to polynomial functions (Ellis and Post 2004, Stenseth et al. 2004, Nevoux et al. 2008). However, regardless of life history complexity or functional response shape, Jensen’s Inequality must be in play if there is any nonlinear fitness-related vital rate at any life history stage. If so, the effect should propagate through the entire population dynamic, influence fitness, and drive apart fitness estimates in the different architectures of environmental fluctuations as variability increases. What would vary between systems is the magnitude of that effect depending on how nonlinear the vital rates are, and how sensitive fitness is to those vital rates. Another open question is if interactions between multiple nonlinear vital rates would amplify or dampen the effect of the mechanisms, to exacerbate or lessen fitness differences between environmental fluctuation architectures.

Perturbation events or periods of harsh conditions are often stark enough to justify focusing on the discrete intervals between them, as we did here, and as is common in disturbance ecology. However, the amplitude, or magnitude, of a continuous environmental variable can itself fluctuate. A prominent example is temperature variability, of particular interest in the context of climate change (Post and Stenseth 1999, AghaKouchak et al. 2020). In the modeling framework we present, variable amplitude can be simulated by allowing the magnitudes of subsequent events to vary randomly or non-randomly, for example in the way events incur mortality or influence the nonlinear vital rate response function following each event. Such analyses might bridge the distinct modelling approaches that either focus on refractory periods between events such as temperature extremes, and those that focus on continuous fluctuations in such variables.

Our general conclusion is that careful and appropriate treatments of the generating processes of environmental fluctuations matter for estimating demographic fitness. In conjunction, while nonlinear change in vital rates within seasons or disturbance phases might be arduous to measure given the temporal sampling required, we argue that more often than not linearity is an assumption made for convenience, not for truthfulness. At least slight nonlinearity is likely a dominant feature of many vital rates, and even if relatively weak, nonlinearity should unlock the synergistic role of the two mechanisms discussed here to drive differences in fitness estimates depending on the architecture of an environment’s temporal fluctuations. Such analyses might significantly expand—as moving from constant to stochastic assumptions did (Tuljapurkar et al. 2009)—our understanding of evolutionary demography so far largely based on stochastic assumptions.

## References

1. AghaKouchak, A. et al. 2020. Climate Extremes and Compound Hazards in a Warming World. - Annu. Rev. Earth Planet. Sci. 48: 519–548.

2. Altizer, S. et al. 2006. Seasonality and the dynamics of infectious diseases. - Ecol. Lett. 9: 467– 484.

3. Boyce, M. S. et al. 2006. Demography in an increasingly variable world. - Trends Ecol. Evol. 21: 141–148.

4. Brown, C. J. et al. 2011. Quantitative approaches in climate change ecology. - Glob. Change Biol. 17: 3697–3713.

5. Bürkli, A. and Jokela, J. 2017. Increase in multiple paternity across the reproductive lifespan in a sperm-storing, hermaphroditic freshwater snail. - Mol. Ecol. 26: 5264–5278.

6. Caswell, H. 2001. Matrix population models: Construction, analysis, and interpretation. 2nd edn sinauer associates. - Inc Sunderland MA in press.

7. Caswell, H. 2008. Perturbation analysis of nonlinear matrix population models. - Demogr. Res. 18: 59–116.

8. Caswell, H. and Trevisan, M. C. 1994. Sensitivity Analysis of Periodic Matrix Models. - Ecology 75: 1299–1303.

9. Caswell, H. and Kaye, T. N. 2001. Stochastic demography and conservation of an endangered perennial plant (Lomatium bradshawii) in a dynamic fire regime. - In: Advances in Ecological Research. Academic Press, pp. 1–51.

10. Chatfield, C. 2003. The Analysis of Time Series: An Introduction, Sixth Edition. - Chapman and Hall/CRC.

11. Chevin, L.-M. et al. 2017. Stochastic Evolutionary Demography under a Fluctuating Optimum Phenotype. - Am. Nat. 190: 786–802.

12. Childs, D. Z. et al. 2010. Evolutionary bet-hedging in the real world: empirical evidence and challenges revealed by plants. - Proc. R. Soc. B Biol. Sci. 277: 3055–3064.

13. Cohen, D. 1966. Optimizing reproduction in a randomly varying environment. - J. Theor. Biol. 12: 119–129.

14. Cole, L. C. 1954. The Population Consequences of Life History Phenomena. - Q. Rev. Biol. 29: 103–137.

15. Crewe, P. et al. 2018. Defining fitness in an uncertain world. - J. Math. Biol. 76: 1059–1099.

16. Cuddington, K. and Hastings, A. 2016. Autocorrelated environmental variation and the establishment of invasive species. - Oikos 125: 1027–1034.

17. Davison, R. et al. 2019. Stochastic effects contribute to population fitness differences. - Ecol. Model. 408: 108760.

18. Denny, M. W. et al. 2009. On the prediction of extreme ecological events. - Ecol. Monogr. 79: 397–421.

19. Drake, J. M. 2005. Population effects of increased climate variation. - Proc. R. Soc. B Biol. Sci. 272: 1823–1827.

20. Dunn, P. O. and Møller, A. P. 2014. Changes in breeding phenology and population size of birds. - J. Anim. Ecol. 83: 729–739.

21. Easterling, D. R. et al. 2000. Climate Extremes: Observations, Modeling, and Impacts. - Science 289: 2068–2074.

22. Ehrlén, J. et al. 2016. Advancing environmentally explicit structured population models of plants. - J. Ecol. 104: 292–305.

23. Ellis, A. M. and Post, E. 2004. Population response to climate change: linear vs. non-linear modeling approaches. - BMC Ecol. 4: 2.

24. Emery, N. C. and La Rosa, R. J. 2019. The Effects of Temporal Variation on Fitness, Functional Traits, and Species Distribution Patterns. - Integr. Comp. Biol. 59: 503–516.

25. Engen, S. et al. 2009. Reproductive Value and the Stochastic Demography of Age-Structured Populations. - Am. Nat. 174: 795–804.

26. Engen, S. et al. 2013. Estimating the effect of temporally autocorrelated environments on the demography of density-independent age-structured populations. - Methods Ecol. Evol. 4: 573–584.

27. Felmy, A. et al. 2021. Ancestral ecological regime shapes reaction to food limitation in the Least Killifish, Heterandria formosa. - Ecol. Evol. 11: 6391–6405.

28. Fieberg, J. and Ellner, S. P. 2001. Stochastic matrix models for conservation and management: a comparative review of methods. - Ecol. Lett. 4: 244–266.

29. Fisher, R. A. 1958. The genetical theory of natural selection. - Dover.

30. Gadgil, M. and Bossert, W. H. 1970. Life Historical Consequences of Natural Selection. - Am. Nat. 104: 1–24.

31. Gaillard, J.-M. et al. 1998. Population dynamics of large herbivores: variable recruitment with constant adult survival. - Trends Ecol. Evol. 13: 58–63.

32. Gaillard, J.-M. et al. 2000. Temporal Variation in Fitness Components and Population Dynamics of Large Herbivores. - Annu. Rev. Ecol. Syst. 31: 367–393.

33. Gonzalez-Andujar, J. L., et al. 2006. Population Cycles Produced by Delayed Density Dependence in an Annual Plant. - Am. Nat. 168: 318–322.

34. Gremer, J. R. and Venable, D. L. 2014. Bet hedging in desert winter annual plants: optimal germination strategies in a variable environment. - Ecol. Lett. 17: 380–387.

35. Haccou, P. and Vatutin, V. 2003. Establishment success and extinction risk in autocorrelated environments. - Theor. Popul. Biol. 64: 303–314.

36. Hacket-Pain, A. and Bogdziewicz, M. 2021. Climate change and plant reproduction: trends and drivers of mast seeding change. - Philos. Trans. R. Soc. B Biol. Sci. 376: 20200379.

37. Halley, J. M. 2005. Comparing aquatic and terrestrial variability: at what scale do ecologists communicate? - Mar. Ecol. Prog. Ser. 304: 274–280.

38. Halley, J. M. and Kunin, W. E. 1999. Extinction Risk and the 1/f Family of Noise Models. - Theor. Popul. Biol. 56: 215–230.

39. Halley, J. M. et al. 2018. How survival curves affect populations’ vulnerability to climate change. - PLOS ONE 13: e0203124.

40. Hayford, H. A. et al. 2018. Radio tracking detects behavioral thermoregulation at a snail’s pace. - J. Exp. Mar. Biol. Ecol. 499: 17–25.

41. Higgins, S. I. et al. 2000. Fire, resprouting and variability: a recipe for grass–tree coexistence in savanna. - J. Ecol. 88: 213–229.

42. Inouye, D. W. 2008. Effects of Climate Change on Phenology, Frost Damage, and Floral Abundance of Montane Wildflowers. - Ecology 89: 353–362.

43. IPCC 2012. Managing the risks of extreme events and disasters to advance climate change adaptation: special report of the Intergovernmental Panel on Climate Change (CB Field, Ed.). - Cambridge University Press.

44. Jensen, J. L. W. V. 1906. Sur les fonctions convexes et les inégalités entre les valeurs moyennes. - Acta Math. 30: 175–193.

45. Jentsch, A. et al. 2007. A new generation of climate-change experiments: events, not trends. - Front. Ecol. Environ. 5: 365–374.

46. Keyfitz, N. 1968. Introduction to the Mathematics of Population.

47. Lande, R. and Orzack, S. H. 1988. Extinction dynamics of age-structured populations in a fluctuating environment. - Proc. Natl. Acad. Sci. 85: 7418–7421.

48. Lande, R. et al. 2003. Stochastic Population Dynamics in Ecology and Conservation. - Oxford University Press.

49. Lande, R. et al. 2017. Evolution of stochastic demography with life history tradeoffs in density-dependent age-structured populations. - Proc. Natl. Acad. Sci. 114: 11582–11590.

50. Lawson, C. R. et al. 2015. Environmental variation and population responses to global change. - Ecol. Lett. 18: 724–736.

51. Lawton, J. H. 1988. More time means more variation. - Nature 334: 563–563.

52. Levins, R. 1968. Evolution in changing environments Princeton University Press. - Princet. N. J. in press.

53. Linderholm, H. W. 2006. Growing season changes in the last century. - Agric. For. Meteorol. 137: 1–14.

54. Lytle, D. A. 2001. Disturbance Regimes and Life-History Evolution. - Am. Nat. 157: 525–536.

55. Lytle, D. A. 2002. Flash Floods and Aquatic Insect Life-History Evolution: Evaluation of Multiple Models. - Ecology 83: 370–385.

56. Lytle, D. A. et al. 2008. Evolution of aquatic insect behaviours across a gradient of disturbance predictability. - Proc. R. Soc. B Biol. Sci. 275: 453–462.

57. MacArthur, R. H. 1962. Some generalized theorems of natural selection. - Proc. Natl. Acad. Sci. U. S. A. 48: 1893.

58. Marlon, J. R. et al. 2009. Wildfire responses to abrupt climate change in North America. - Proc. Natl. Acad. Sci. 106: 2519–2524.

59. McAdam, A. G. and Boutin, S. 2003. Variation in Viability Selection Among Cohorts of Juvenile Red Squirrels (tamiasciurus Hudsonicus). - Evolution 57: 1689–1697.

60. Metcalf, C. j. e and Koons, D. n 2007. Environmental uncertainty, autocorrelation and the evolution of survival. - Proc. R. Soc. B Biol. Sci. 274: 2153–2160.

61. Morris, W. F. et al. 2006. Sensitivity of the population growth rate to demographic variability within and between phases of the disturbance cycle. - Ecol. Lett. 9: 1331–1341.

62. Morris, W. F. et al. 2008. Longevity Can Buffer Plant and Animal Populations Against Changing Climatic Variability. - Ecology 89: 19–25.

63. Nevoux, M. et al. 2008. Nonlinear impact of climate on survival in a migratory white stork population. - J. Anim. Ecol. 77: 1143–1152.

64. Noble, J. D. et al. 2019. Foraging strategies of individual silky pocket mice over a boom–bust cycle in a stochastic dryland ecosystem. - Oecologia 190: 569–578.

65. Nouvellet, P. et al. 2012. Noisy clocks and silent sunrises: measurement methods of daily activity pattern. - J. Zool. 286: 179–184.

66. Park, J. S. 2019. Cyclical environments drive variation in life-history strategies: a general theory of cyclical phenology. - Proc. R. Soc. B 286: 20190214.

67. Park, J. S. and Wootton, J. T. 2021. Slower environmental cycles maintain greater life-history variation within populations. - Ecol. Lett. 24: 2452–2463.

68. Pélisson, P.-F. et al. 2012. Contrasted breeding strategies in four sympatric sibling insect species: when a proovigenic and capital breeder copes with a stochastic environment. - Funct. Ecol. 26: 198–206.

69. Pike, N. et al. 2004. The effect of autocorrelation in environmental variability on the persistence of populations: an experimental test. - Proc. R. Soc. Lond. B Biol. Sci. 271: 2143–2148.

70. Post, E. and Stenseth, N. Chr. 1999. Climatic Variability, Plant Phenology, and Northern Ungulates. - Ecology 80: 1322–1339.

71. Priestley, M. B. 1981. Spectral analysis and time series: probability and mathematical statistics. - Academic Press.

72. Pyper, B. J. and Peterman, R. M. 1998. Comparison of methods to account for autocorrelation in correlation analyses of fish data. - Can. J. Fish. Aquat. Sci. 55: 2127–2140.

73. Reuman, D. C. et al. 2008. Colour of environmental noise affects the nonlinear dynamics of cycling, stage-structured populations. - Ecol. Lett. 11: 820–830.

74. Ruel, J. J. and Ayres, M. P. 1999. Jensen’s inequality predicts effects of environmental variation. - Trends Ecol. Evol. 14: 361–366.

75. Sabo, J. L. and Post, D. M. 2008. Quantifying Periodic, Stochastic, and Catastrophic Environmental Variation. - Ecol. Monogr. 78: 19–40.

76. Schaffer, W. M. 1974. Optimal Reproductive Effort in Fluctuating Environments. - Am. Nat. 108: 783–790.

77. Schreiber, S. J. and Moore, J. L. 2018. The structured demography of open populations in fluctuating environments. - Methods Ecol. Evol. 9: 1569–1580.

78. Shriver, R. K. 2016. Quantifying how short-term environmental variation leads to long-term demographic responses to climate change. - J. Ecol. 104: 65–78.

79. Sletvold, N. et al. 2013. Climate warming alters effects of management on population viability of threatened species: results from a 30-year experimental study on a rare orchid. - Glob. Change Biol. 19: 2729–2738.

80. Staver, A. C. et al. 2009. Browsing and fire interact to suppress tree density in an African savanna. - Ecol. Appl. 19: 1909–1919.

81. Stearns, S. C. 1976. Life-History Tactics: A Review of the Ideas. - Q. Rev. Biol. 51: 3–47.

82. Steele, J. H. 1985. A comparison of terrestrial and marine ecological systems. - Nature 313: 355–358.

83. Stenseth, N. Chr., et al. 2004. Modelling non–additive and nonlinear signals from climatic noise in ecological time series: Soay sheep as an example. - Proc. R. Soc. Lond. B Biol. Sci. 271: 1985–1993.

84. Taylor, C. et al. 2014. Nonlinear Effects of Stand Age on Fire Severity. - Conserv. Lett. 7: 355– 370.

85. Thornton, P. K. et al. 2014. Climate variability and vulnerability to climate change: a review. - Glob. Change Biol. 20: 3313–3328.

86. Tuljapurkar, S. 1985. Population dynamics in variable environments. VI. Cyclical environments. - Theor. Popul. Biol. 28: 1–17.

87. Tuljapurkar, S. 1990. Population Dynamics in Variable Environments. - Springer-Verlag.

88. Tuljapurkar, S. and Haridas, C. V. 2006. Temporal autocorrelation and stochastic population growth. - Ecol. Lett. 9: 327–337.

89. Tuljapurkar, S. et al. 2003. The Many Growth Rates and Elasticities of Populations in Random Environments. - Am. Nat. 162: 489–502.

90. Tuljapurkar, S. et al. 2009. From stochastic environments to life histories and back. - Philos. Trans. R. Soc. B Biol. Sci. 364: 1499–1509.

91. van de Pol, M. et al. 2011. Poor environmental tracking can make extinction risk insensitive to the colour of environmental noise. - Proc. R. Soc. B Biol. Sci. 278: 3713–3722.

92. Vasseur, D. A. and Yodzis, P. 2004. The Color of Environmental Noise. - Ecology 85: 1146– 1152.

93. Venable, D. L. 2007. Bet Hedging in a Guild of Desert Annuals. - Ecology 88: 1086–1090.

94. Walther, G.-R. et al. 2002. Ecological responses to recent climate change. - Nature 416: 389–395.

95. Westerling, A. L. et al. 2006. Warming and Earlier Spring Increase Western U.S. Forest Wildfire Activity. - Science 313: 940–943.

