## Supplementary material for "Why the architecture of environmental fluctuation matters for fitness": Fig. S

Supplementary Figures

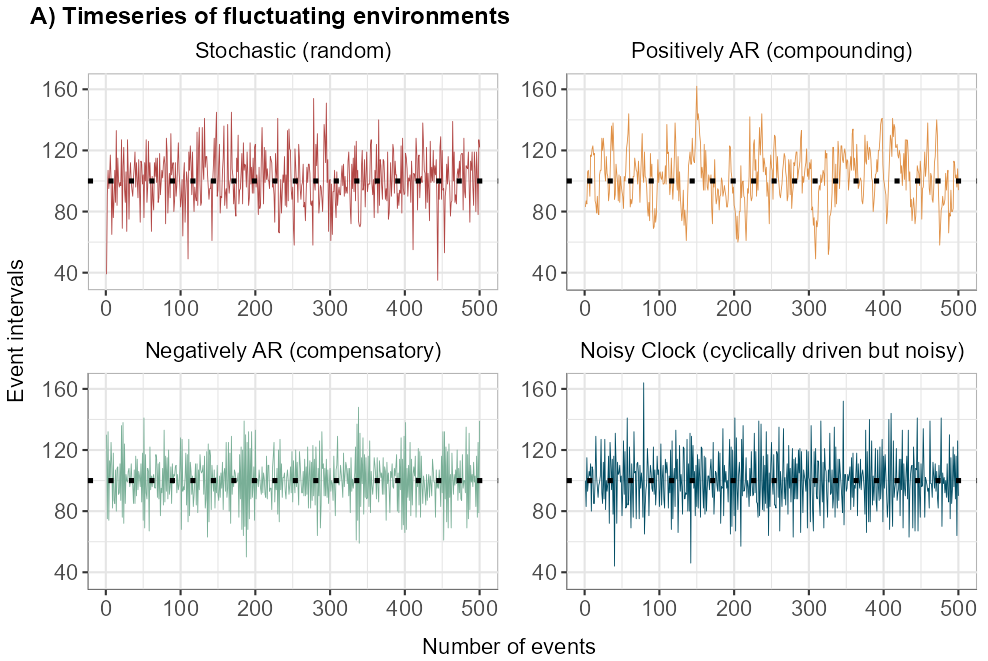

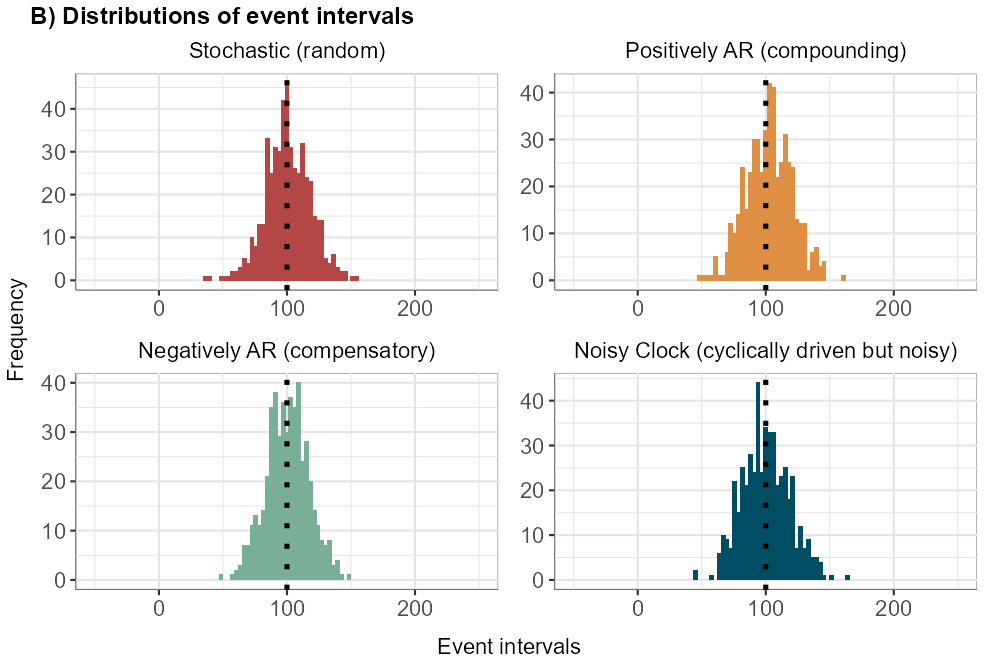

**Figure S1.** A) Timeseries of event intervals produced by the four generating functions. B) The frequency distributions of simulated event intervals are equivalent normal distributions (Table 1 in main text). Dotted line shows imposed mean interval, in this case 100 time steps (days).

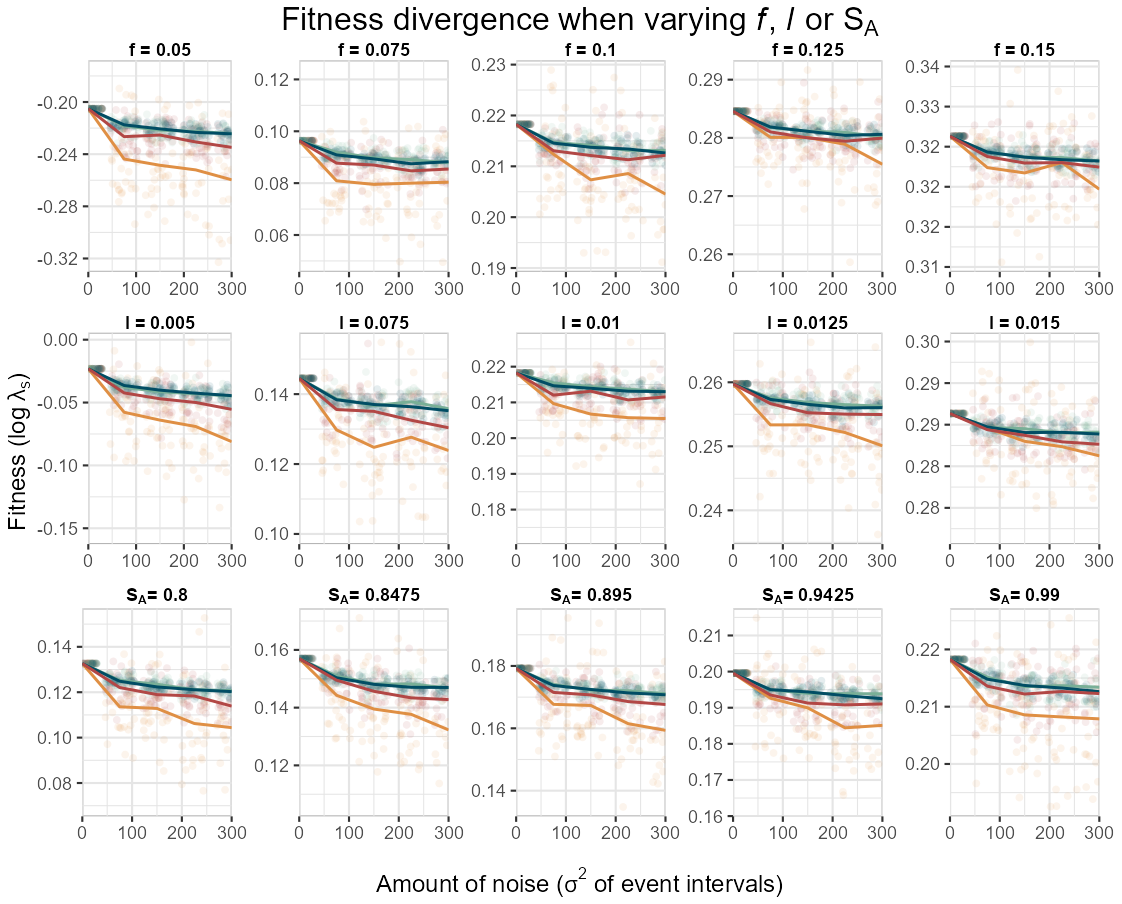

**Figure S2.** Varying the fixed values of daily reproductive rate ($f$), maturation rate ($l$), or adult survival ($S_{A}$) at disturbance in the demographic model does not alter the height order nor the qualitative shapes of the fitness (${log}_{e}\lambda_{s}$) curves among the four fluctuation architectures as amount of noise increases. The overal heights of all curves (y-axis) increase as any of the 3 variables increase (all 3 rows, left to right) because the parameters all contribute to faster growth of the population.
